# A comparison of short- and long-read whole genome sequencing for microbial pathogen epidemiology

**DOI:** 10.1101/2025.02.17.638699

**Authors:** Andrea M. Schiffer, Arafat Rahman, Wendy Sutton, Melodie L. Putnam, Alexandra J. Weisberg

## Abstract

Whole genome sequencing provides the highest resolution for characterizing pathogen evolution, epidemiology, and diagnostics. Genome assemblies contain information on the identity and potential phenotypes of a pathogen. Likewise, variant calling can inform on transmission patterns and evolutionary relationships. Recent improvements in Oxford Nanopore long-read sequencing have made its use attractive for genomic epidemiology. However, the accuracy and optimal strategy for analysis of Nanopore reads remains to be determined. We compared the use of Illumina short reads and Oxford Nanopore long reads for genome assembly and variant calling of phytopathogenic bacteria. We generated short- and long-read datasets for diverse phytopathogenic *Agrobacterium* strains. We then analyzed these data using multiple pipelines designed for either short or long reads and compared the results. We found that assemblies made from long reads were more complete than those made from short-read data and contained few sequence errors. Variant calling pipelines differed in their ability to accurately call variants and infer genotypes from long reads. Results suggest that computationally fragmenting long reads can improve the accuracy of variant calling in population-level studies. Using fragmented long reads, pipelines designed for short reads were more accurate at recovering genotypes than pipelines designed for long reads. Further, short- and long-read datasets can be analyzed together with the same pipelines. These findings show that Oxford Nanopore sequencing is accurate and can be sufficient for microbial pathogen genomics and epidemiology. Ultimately, this enhances the ability of researchers and clinicians to understand and mitigate the spread of pathogens.

**Importance:** Genome assembly and variant calling are important steps in microbial population studies and epidemiology. Most variant calling and genotyping pipelines are designed for Illumina short sequencing reads. Oxford Nanopore Technology long-read sequencing results in more complete genome assemblies but has historically been of lower quality. Here, we show that Nanopore long reads are now of sufficient quality for bacterial whole genome assembly and epidemiology. We benchmarked the accuracy of multiple variant-calling pipelines with short and long reads. Using an optimized variant calling approach, variant calls and genotypes inferred from long reads are as accurate as those inferred from short reads. Importantly, we found that gold-standard variant calling pipelines designed for short reads are also accurate with long reads when long reads are first fragmented into shorter sequences. This finding allows researchers to incorporate the advantages of Nanopore sequencing for genome assembly, while maintaining high accuracy for epidemiology and population analysis.

## Introduction

Whole genome sequencing (WGS) has driven major advances in understanding plant and clinical pathogen evolution, epidemiology, and diagnostics (1–4). Whole genome sequences can be used to infer host range, antimicrobial resistance, and other characteristics of microbial plant and clinical pathogens. WGS data are advantageous because they simultaneously provide the highest resolution for epidemiological analysis and the characterization of pathogen transmission patterns. Recent technological developments have made it possible for plant and clinical diagnostic clinics to incorporate these tools into their arsenal. These data have already informed on emerging pathogens, such as the recent outbreak of multidrug-resistant *Pseudomonas aeruginosa* in contaminated eye drops, and an introduction of *Xanthomonas hortorum* pv. *pelargonii* into the U.S. on geranium (5, 6).

Traditionally, short-read Illumina sequencing has been considered the gold standard for whole genome sequencing. Illumina sequencing produces reads with high accuracy, which can be used to assemble genomes with high consensus sequence confidence or for whole genome variant calling (7). However, Illumina reads are at most 300 bp in length, which limits their use in assembling repetitive regions of genomes. Therefore, most genome assemblies made from short reads are fragmented into multiple pieces, called contigs. Chromosome structure or the presence of mobile genetic elements such as plasmids can be difficult to infer from short read assemblies.

The introduction of long-read sequencing aimed to resolve many of the issues inherent to short-read sequencing. Oxford Nanopore sequencing has the advantage of producing very long reads, up to a megabase in length, and on sequencers that are affordable to most labs. However, previous studies found that the quality of Nanopore reads was not high enough to be used on their own (8, 9). Nanopore reads also contained homopolymer repeat errors that can result in indels in assemblies (8, 10). Therefore, short-read sequencing data was often necessary for polishing long-read assemblies (11). However, recent advances in Oxford Nanopore chemistry and basecalling have changed the landscape of long-read sequencing. Assemblies produced from Nanopore long reads are now nearly perfect for many organisms (12).

Epidemiology and pathogen transmission can be characterized to the highest resolution using whole genome variant calling. Variant calling pipelines typically use the alignment of reads to a common reference genome to identify single nucleotide polymorphisms (SNPs) for each strain (13). Strains differing by less than a determined threshold, often 10-15 SNPs, are then grouped into genotypes (14). Mapping host or location information onto genotypes can then be used to characterize pathogen transmission. Short-read data has long been used for variant calling, and mature tools are available for this analysis (15–17). However, the historically poor quality of long reads has meant that their value and use in variant calling is not well characterized. Tools for variant calling with Nanopore reads are now available (18).

In this study, we assessed the quality of long reads in genome assembly and variant calling for population-level genotyping analysis. We compared genome assemblies made from either short or long reads. We also compared different strategies for variant calling using tools designed for either Illumina or Nanopore reads. We analyzed the consistency of variant calls produced from short and long reads of the same strain. We also assessed the ability of each pipeline to correctly group short and long read data from the same strain, or pairs of strains known to be identical, into the same genotype. Finally, we highlight the potential benefits that plant and medical clinics can derive from adding whole genome sequencing to their pathogen diagnostic testing.

## Results

To test the viability of long-read sequencing for bacterial epidemiology, we analyzed 116 putative *Agrobacterium* strains isolated from plant samples submitted to the OSU Plant Clinic. We also included strains from the Larry Moore and Thomas Burr culture collections, and from the California Department of Food and Agriculture (CDFA) collection. These 116 strains were selected to maximize diversity in plant host, location, and year of isolation (**Table S1**). We generated Illumina short-read data for all strains, and Oxford Nanopore long-reads for a random subset of 35 strains and analyzed them independently using multiple established pipelines. The 35 paired datasets allowed us to compare standard pipelines using short reads with assembly and variant calling pipelines using long reads. The accuracy and completeness of each approach were assessed using multiple metrics.

We first assessed the quality of short- and long-read datasets. The average read length of Nanopore reads was 6,835 bp (mean N50 read length 13,385 bp; **Table S2**). Average coverage for Illumina read datasets varied from 22X to 106X (mean 52X coverage) and average coverage for Nanopore read datasets varied from 17X to 173X (mean 61X coverage; **Table S3**). During this study, multiple Nanopore basecalling models were released, so reads were rebasecalled and retested in the pipeline. We compared the “SUP” versions of production model v4.2.0, a beta-release bacterial methylation-aware model (hereafter “bacmethyl”), and the latest production model v5.0.0. The quality of reads was consistent or improved with each new model, particularly with v5.0.0 (**Table S2**). Nanopore simplex reads basecalled with the v5.0.0 SUP model had an average quality score of Q19.12, or a per-read accuracy of ∼98.7%.

### Assemblies made with Nanopore data reveal complete chromosome structure

Genomes were assembled from either short- or long-read data, or a hybrid approach combining both, and then compared to assess completeness and accuracy (19–21). As expected, genomes assembled from long-read data or hybrid data were more complete and had higher N50 values than those assembled from short reads (**Figure 1A and B; Table S3**). The average N50 was 483,620 bp for short-read SPAdes assemblies and 3,784,004 bp for long-read Flye assemblies. Short-read datasets assembled into an average of 55 contigs (**Table S3**). In contrast, nearly all long-read assemblies contained structurally complete chromosomes and plasmids, if present (**Table S3**). Total assembly size was comparable between all assemblies, but Flye and hybrid assemblies were generally slightly larger (**Table S3**).

**Figure 1:**
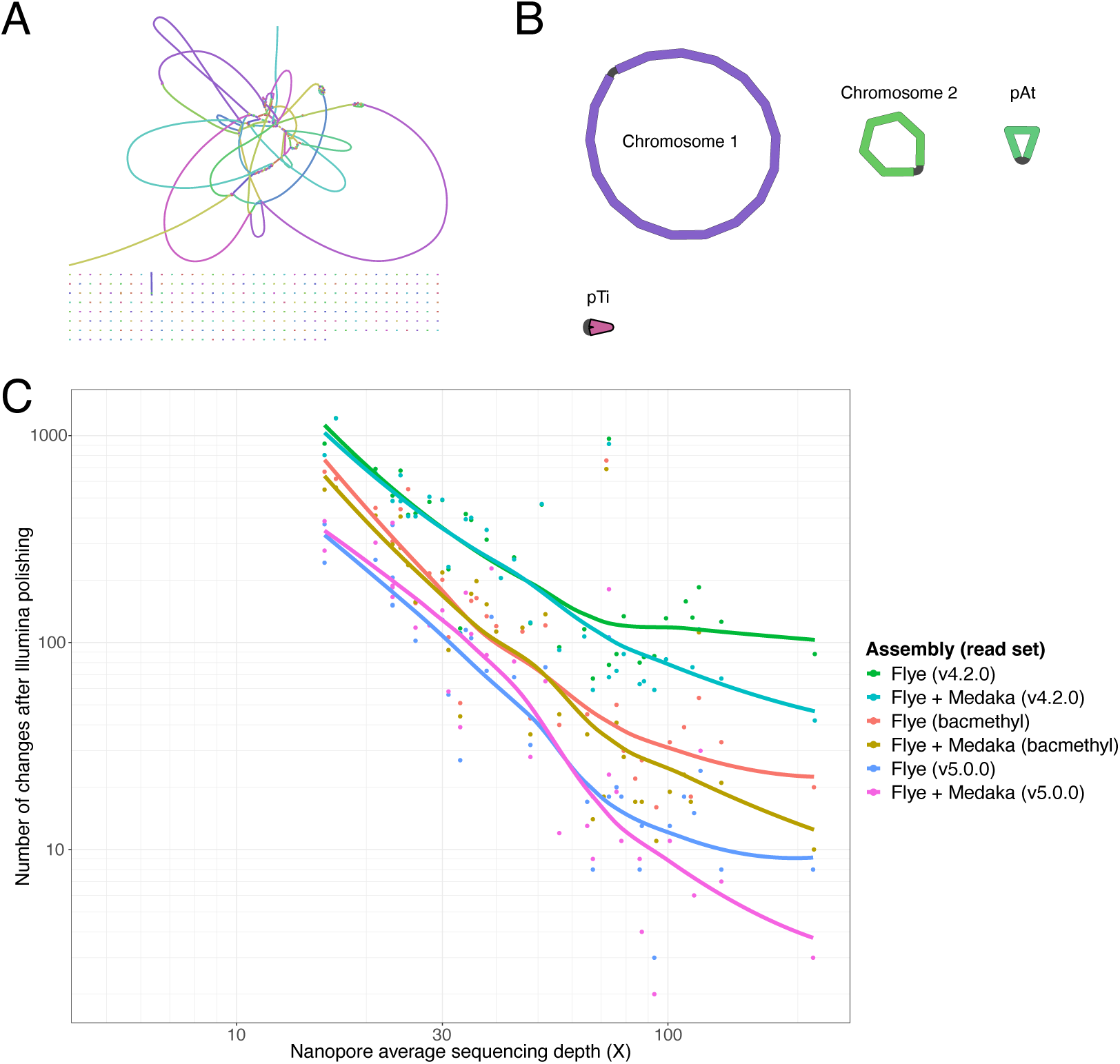
Nanopore-only assemblies with sufficient depth of sequencing are complete and have few errors. Assembly graphs of strain CG154 assembled from **A.** Illumina short reads or **B.** Nanopore long reads. **C.** Assemblies of sequenced strains plotted as points on a graph of long read sequencing depth (X) by the number of changes made to that assembly following Polypolish polishing with Illumina short reads. The long-read assembly method (Flye or Flye with Medaka polishing) and basecalling model version (v4.2.0, beta methylation-aware bacteria model [bacmethyl], or v5.0.0) is indicated by color. A best-fit line was plotted for each assembly method and model combination using locally estimated scatterplot smoothing.

### Improved basecalling reduces assembly errors in long read-only assemblies

Genomes assembled from long reads have historically contained many single nucleotide polymorphism (SNP) and small insertion/deletion (indel) errors (22). However, recent improvements in Nanopore basecalling models have reduced this error rate. We assessed the number of assembly errors in each long-read assembly by polishing with high quality Illumina short reads and counting the number of changes made (11).

Assemblies made from reads basecalled with the most recent model (v5.0.0) had the fewest assembly errors (**Table S4**). Long-read assemblies with at least 50x coverage had on average 23 SNP errors, with as few as 3 errors. The number of errors in long-read assemblies declined with increasing sequencing depth of long reads (**Figure 1C**). Strains with less than 30X long-read coverage had many errors, while strains with >30X sequencing depth had fewer errors (mean 43 errors). Above 80X long-read coverage, the number of errors did not dramatically decrease as sequencing depth increased. We also tested the effectiveness of polishing long-read assemblies with long reads using Medaka (23). Medaka polishing slightly reduced the number of assembly errors when long-read datasets were >80X coverage (**Figure 1C; Table S4**).

We noticed an unusually high number of errors in the assembly of strain CG160, despite this sample having >100X long-read coverage. We identified low levels of contamination in the short and long reads of this sample. This contamination primarily affected the assembly of one plasmid, which contained nearly all assembly errors. Manual examination of read alignments revealed that errors were in conserved plasmid replication and conjugation loci. After filtering reads that mapped to the contaminants, a new assembly made from the filtered reads reduced the errors from 469 to 24 (**Table S4**).

We next identified motifs in long-read assemblies associated with errors. Previous studies found that errors in Nanopore reads occur more frequently near methylated nucleotides (24). Bacterial DNA is often methylated at specific sequence motifs (25). The motif “GANTC” or “GATC” is methylated as part of cell cycle regulation in many Alpha- and Gammaproteobacteria, respectively (26–28). Sequence motifs targeted by restriction-modification (R-M) defense systems are also methylated (29, 30). We characterized the sequence context around errors in long-read assemblies and identified conserved motifs (**Table S5**). Most assembly SNP errors are near sites likely to be targeted for methylation. Nearly all identified motifs are either known to be methylated in agrobacteria, such as “GANTC”, or are short palindromes that resemble R-M system methylation targets, such as “CTGCAG”. Overall, assembly errors were reduced in datasets made from basecalling models that account for methylation (bacmethyl, v5.0.0) relative to models that do not (v4.2.0; **Figure 1C; Table S4**). Most improvements in assembly error rates are in sites with predicted methylated motifs (**Table S5**). Other assembly errors were identified as small indels or homopolymer errors, though these also were reduced with more recent basecalling models (**Table S4**).

We compared gene annotations of SPAdes short-read assemblies, unpolished and short-read polished Flye long-read assemblies, and Unicycler hybrid assemblies to characterize the quality of gene calls. Flye assemblies, both polished and unpolished, consistently had slightly greater numbers of annotated gene features than SPAdes assemblies (**Figure S1**). There were few or no differences in the number of genes called between polished and unpolished Flye assemblies (**Table S3; Figure S1**). BUSCO scores were comparable across all read-type and assembly methods (**Table S3; Figure S2**).

### SNP calling pipelines vary in accuracy when used with long-read data

We next compared short and long reads for whole genome variant calling using a variety of pipelines. We tested four SNP calling pipelines with short reads and v5.0.0 long reads. Three of the pipelines, GATK, Graphtyper, and Bcftools, were originally designed for Illumina short-read data (15–17). The other tool, Clair3, is the only one specifically designed for variant calling of Nanopore reads (18). We previously found Graphtyper to be the most consistently accurate for inferring clonal genotypes of bacterial pathogens from short-read data (31, 32). GATK was also accurate if datasets were first partitioned into below species-level groups. While neither tool was designed for long reads, we tested these pipelines with our short- and long-read datasets. We then assessed the consistency of variant calls across short and long read datasets.

The analyzed strains represent genetically diverse agrobacteria (**Figure S3**). The newly sequenced strains belong to multiple species and represent each of the major pathogen lineages within the agrobacteria/rhizobia complex. Variant calling was initially performed within species-level groups (ANI >95%) and with the quality filtering thresholds recommended by each pipeline. For each strain, short and long reads were analyzed separately in each pipeline, along with strains from NCBI. We then compared the number of SNPs to the reference inferred for each strain with either short- or long-read data (**Figure 2A; Table S6**).

**Figure 2:**
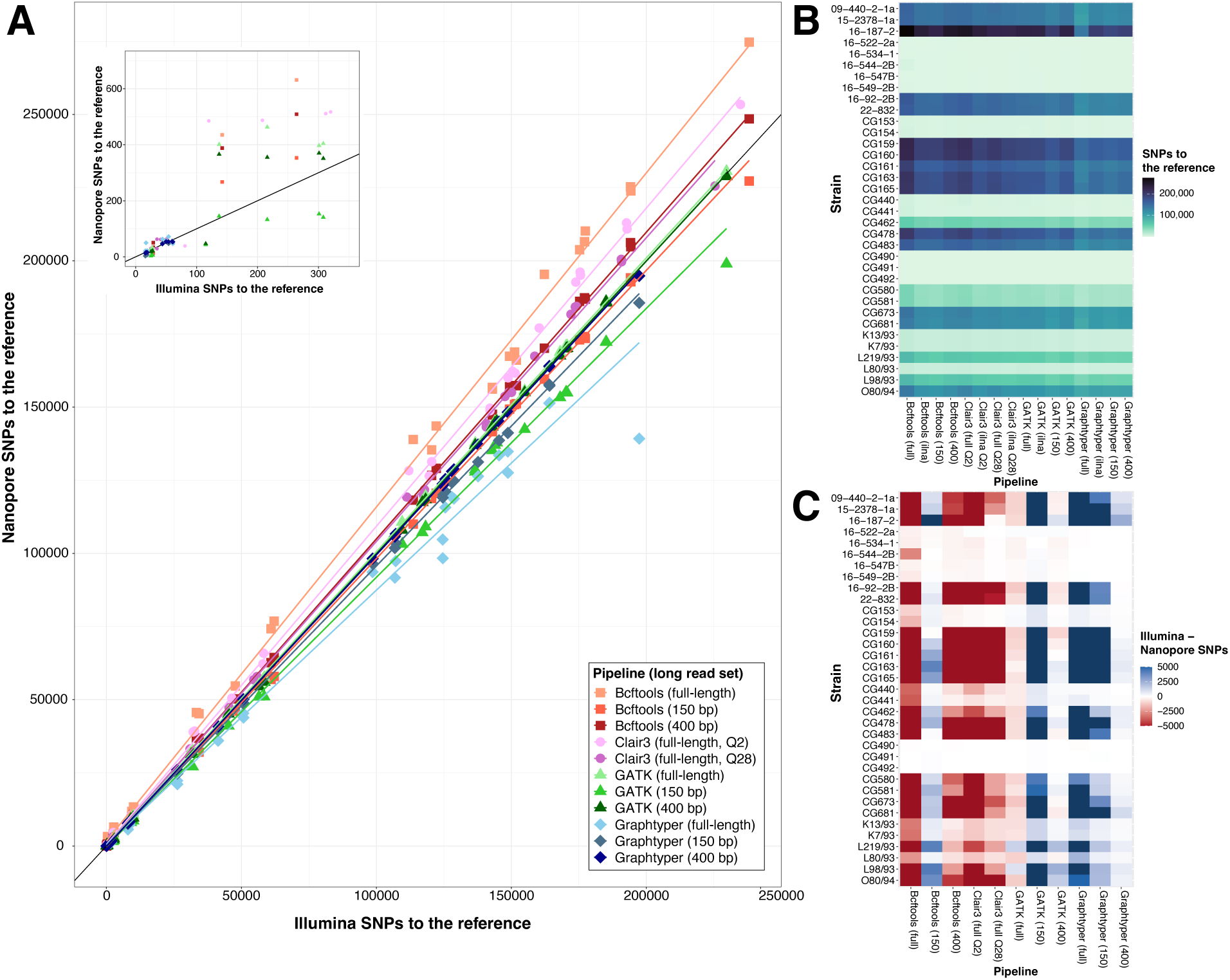
Comparison of variant calling pipelines. **A.** Plot of SNP calls to the reference for Illumina vs Nanopore reads for each strain and pipeline. Each point represents an individual strain, shape and color indicate variant calling pipeline and long-read dataset type, respectively. Long reads were either full length or fragmented into 150 bp (150) or 400 bp (400) pieces. Quality filtering for Clair3 was performed with either a threshold of QUAL > 2 (Q2) or QUAL > 28 (Q28). The black line represents a predicted 1 to 1 ratio between Illumina SNPs to the reference and Nanopore SNPs to the reference. The embedded graph is zoomed to show the 0 to 350 SNPs region. **B.** A heatmap comparing the number of SNPs called to a reference using different variant calling pipelines and read sets. Each row is a strain, each column is the number of SNPs called for that strain using each pipeline and read type. **C.** A heatmap representing the difference in SNP calls to the reference made with Illumina and Nanopore reads for each strain. Rows are strains, columns are variant calling pipelines and long-read type. Color indicates whether more SNPs were called with Illumina reads (blue) or Nanopore reads (red). Values greater than 5000 SNP differences are colored dark red and dark blue.

Overall, all pipelines called roughly comparable numbers of SNPs to the reference for each strain with short and full-length long reads (**Figure 2B**). However, each pipeline differed in how consistently calls were made with each read type. Bcftools and Clair3 consistently called more SNPs with full-length long reads than with short reads for each strain (**Figure 2A and C**). In contrast, Graphtyper consistently called fewer variants with full-length long reads than with short reads. GATK called similar numbers of variants with either short or full-length long reads. When considering only comparisons where strains are closely related to the reference genome (<300 SNPs), all pipelines produced largely consistent SNP calls using either short-read or full-length long-read data (**Figure 2A inset**).

### Fragmented long reads are comparable to short reads in variant calling pipelines

To more closely approximate variant calling with short reads, we fragmented full-length long reads into either 150 bp or 400 bp sequences, aligned them to reference genomes, and called variants using each pipeline. Fragmentation of long reads led to more consistent SNP calls with short reads for Graphtyper, GATK, and Bcftools, but not for Clair3 (**Figure 2; Tables S7-10**). Variant calling using 400 bp fragments was most consistent with calls for short-read data when using Graphtyper or GATK, while variant calling with 150 bp fragments was most consistent when using Bcftools. Variant calling with Graphtyper or GATK pipelines using 400 bp long-read fragments, Bcftools with 150 bp long-read fragments, and Clair3 with full-length long reads most closely matched short-read SNP calls (**Figure 2A, S4**).

We initially hypothesized that poor read mapping of full-length long reads to distantly related reference genomes was responsible for the under-calling of variants with some pipelines. However, the mapped read depth, breadth, and quality of full-length reads was not significantly worse than for 400 bp fragmented reads across datasets (**Figure S5**). In fact, all mapping statistics were slightly better with full-length reads relative to fragmented reads in nearly all datasets (**Table S11**). Therefore, we hypothesized that pipelines designed for short reads were unable to correctly handle long reads. To test this, we fragmented full-length mapped reads in place into 400 bp fragments, retaining the exact mapping position and quality of each sub-sequence of the full-length read alignment. Variant calls produced by Graphtyper with 400 bp split-in-place long reads were comparable to variant calls for individually mapped 400 bp read fragments (**Table S10**). Variant calls for both split-in-place and independently mapped 400 bp read fragments were more consistent with variant calls for short reads than full-length long reads (**Table S10)**.

### Variant calling with Clair3 is improved by more stringent quality filtering

To infer the accuracy of variant calls with long reads, we calculated precision, recall, and F1 scores for each pipeline. Variant calls produced from short reads with each pipeline, or from short reads with Graphtyper, were used as truth sets. Variant calls produced by each pipeline from either full-length or fragmented long reads were used as the test set. Precision vs. recall curves of variant calls across all strains were used to identify optimal filtering thresholds for each pipeline and read set (**Figure S6-S7**). Using the same pipeline with short reads as the truth set, most pipelines had little variation in precision or recall across thresholds (**Figure S6**). However, Clair3 had a maximized precision and recall at a median threshold of QUAL > ∼28. Therefore, we also included Clair3 with a variant filtering threshold of QUAL > 28 in all further comparisons (**Table S13**). Precision vs. recall curves made using Graphtyper with short reads as the truth set were largely consistent with the analyses using within-pipeline truth sets (**Figure S7**).

### Different long read sets are most accurate with each pipeline

We next compared the overall precision, recall, and F1 scores across pipelines and read sets (**Figure 3**). When using the same pipeline as a truth set, Graphtyper with 400 bp reads had significantly greater precision than all other pipelines except for GATK with 400 bp reads (Kruskal-Wallis test *p*-value <2.2*10^-16^, Dunn’s post-hoc test; **Figure 3; Table S12**). Graphtyper with 400 bp reads also had significantly greater F1 scores than all other pipelines except for GATK with 400 bp reads and Clair3 with full-length reads and a filtering threshold of QUAL > 28 (Kruskal-Wallis test *p*-value <2.6*10^-14^, Dunn’s post-hoc test; **Figure 3**). Recall scores were not significantly different across most pipelines and read sets. Graphtyper had comparable precision with either full-length or 400 bp reads, but significantly worse recall and F1 scores with full-length reads (Kruskal-Wallis test *p*-value <2.2*10^-16^, Dunn’s post-hoc test). Graphtyper with 400 bp long reads had a median precision of 99.2%, median recall of 98.5%, and median F1 of 98.8%. GATK with 400 bp long reads had a median precision of 97.8%, median recall of 95.9%, and median F1 of 96.1%. Clair3 with full-length reads and a QUAL > 28 filtering threshold had a median precision of 95%, median recall of 99.2%, and median F1 of 96.9%.

**Figure 3:**
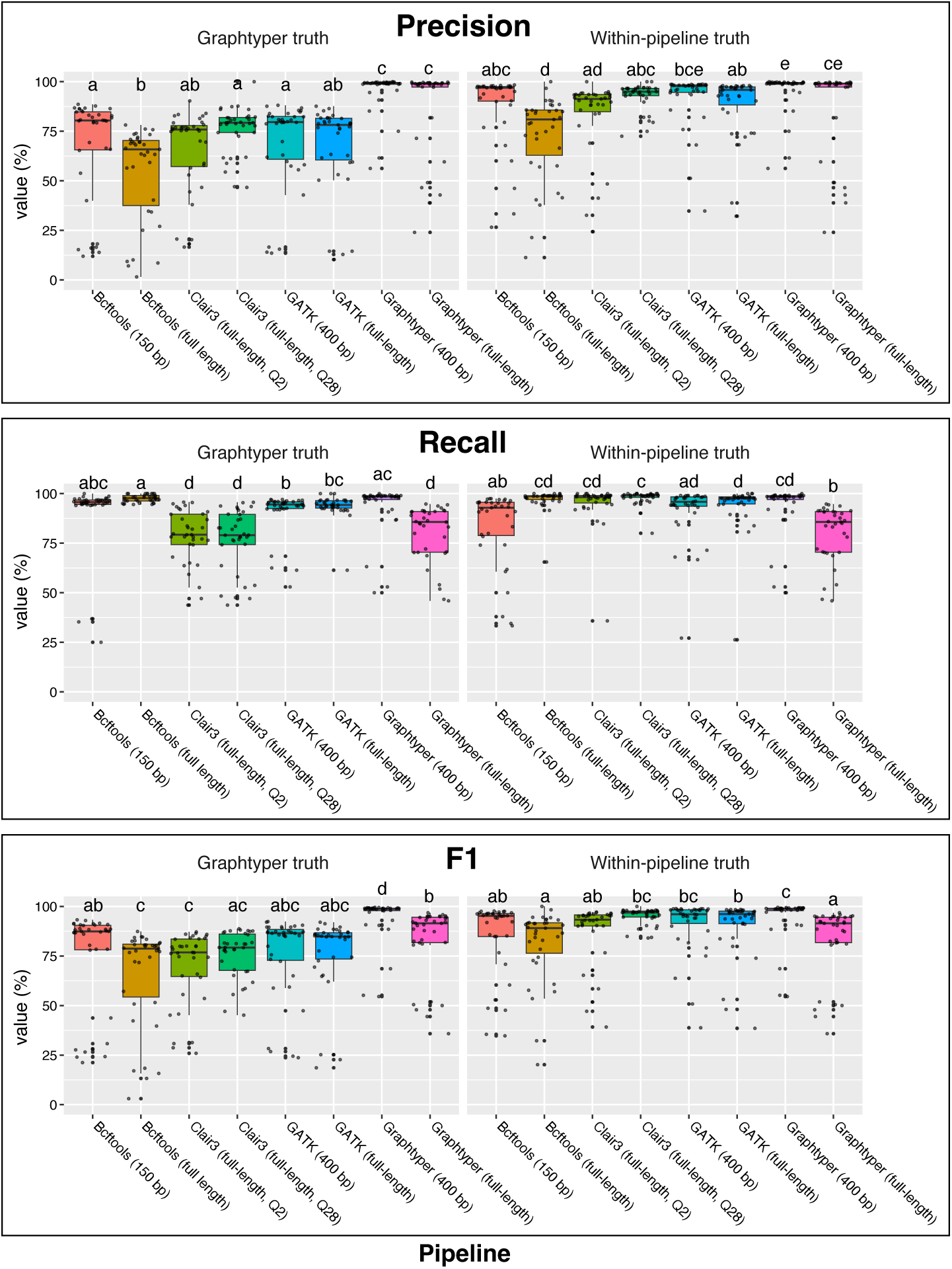
Variant calling accuracy across pipelines. Precision, recall, and F1 scores for per-strain variant calls produced by each pipeline using either full-length or fragmented long reads. Truth sets were either variant calls produced by Graphtyper from Illumina short reads or variant calls produced by the tested pipeline using Illumina short reads. Scores are represented as percentages. Scores for each strain are represented as points and summarized as boxplots colored by dataset. Significance letters from a Dunn’s post-hoc test are represented above each boxplot.

When variant calls produced by Graphtyper using short reads were used as a truth set, Graphtyper with either full-length or 400 bp reads had significantly better precision than all other pipelines (Kruskal-Wallis test *p*-value <2.2*10^-16^; **Figure 3**). Graphtyper with 400 bp reads had significantly better F1 scores than all other pipelines and read sets, including Graphtyper with full-length reads (Kruskal-Wallis test *p*-value <2.2*10^-16^, Dunn’s post-hoc test). Graphtyper with 400 bp reads had significantly better recall than all other pipelines except for Bcftools with 150 bp reads (Kruskal-Wallis test *p*-value <2.2*10^-16^, Dunn’s post-hoc test; **Figure 3**). All other pipelines had similar precision, recall, and F1 scores.

### Long reads are sufficient for accurately inferring genotypes

We next compared pipelines for inferring genotypes from SNP data. Genotyping, identifying sets of genetically identical strains, is important for characterizing outbreaks and understanding pathogen transmission (14, 33). For each pipeline or approach, we characterized whether short- and long-read datasets for the same strain were correctly inferred to be the same genotype in 95% ANI population-level analyses. A threshold of ≤15 relative pairwise SNP differences was used to infer two strains as belonging to the same genotype. Graphtyper with 400 bp long reads correctly paired the most pairs of short and long read datasets into genotypes (**Table S14**). Graphtyper correctly inferred short-read and 400 bp long-read data, fragmented either in-place or pre-mapping, from the same strain into genotypes for 18 and 16 of 35 strains, respectively. Graphtyper with full-length or 150 bp fragments correctly inferred 4 and 14 of 35 same-strain genotypes with short reads, respectively. GATK with full-length reads or any fragmented read type correctly inferred 8 and 7 genotypes, respectively. Clair3 with full-length reads and QUAL > 28 filtering correctly inferred the same genotype for 8 of 35 strains. Clair3 with a filtering threshold of QUAL > 2 correctly inferred 3 of 35 strains, while Bcftools with either full-length or 400 bp reads correctly inferred genotypes for just 2 of 35 strains.

While Graphtyper with 400 bp long-read fragments correctly paired short and long reads into genotypes for most strains, it did not group all. We hypothesized that this may be due to comparing strains to distantly related references. Using a 95% ANI threshold, strains with 5-7 Mb genomes can be grouped with those that differ by ∼100,000 SNPs. Therefore, we tested partitioning the strains into groups using a 99% ANI threshold. We tested Graphtyper and Clair3 with these groupings (**Tables S15, S16**). While some strains could not be analyzed because the 99% ANI threshold grouped them as singletons, Graphtyper correctly grouped short-read and 400 bp long-read datasets for 26 of 28 strains (**Tables S14, S15**). Clair3 with a filtering threshold of QUAL > 28 correctly grouped short-read and full-length long-read datasets for 11 of 28 strains (**Tables S14, S16**).

### Pipelines differ in their ability to correctly genotype known identical strains

We next characterized how consistently each pipeline grouped different strains into genotypes. We identified 20 pairs of strains that are nearly identical (≤15 pairwise SNPs) based on alignment of short reads from one strain to the assembly of the other and calling variants with Graphtyper (**Table S17**). We identified 19 of the same strain pair genotypes when Clair3 and full-length long reads, with QUAL > 2 filtering, was used instead (**Table S17**). However, the final strain pair, K13/93 and K7/93, was found to be nearly identical when a filtering threshold of QUAL > 28 was used (**Table S17**). Therefore, we used these 20 strain pairs as truth set genotypes. We then characterized the ability of each pipeline to correctly genotype these strains with either short or long reads mapped to distant reference genomes in 95% ANI population-level analyses (**Table S18**).

Graphtyper with either short reads or 400 bp fragmented long reads was able to correctly genotype the most known strain pairs, identifying 17 of 20 truth-set genotypes (**Tables S10, S18**). Graphtyper with 150 bp fragments or full-length reads correctly inferred 16 and 3 genotypes, respectively. The next most accurate pipeline was Clair3 with long reads and a filtering threshold of QUAL > 28, which correctly inferred 15 genotypes (**Tables S13, S18**). Clair3 with long reads and QUAL > 2 filtering inferred 11 truth set genotypes (**Tables S8, S18**). GATK correctly inferred 7 truth set genotypes with either full-length or 150 bp fragmented reads (**Tables S9, S18**). Bcftools with any long read dataset correctly inferred 1 truth set genotype (**Table S7, S18**). Bcftools with short reads correctly inferred 9 truth set genotypes. Genotypes were inferred by comparing strains sequenced with the same technology and read type. However, pipelines varied in their ability to correctly infer known genotypes for strains sequenced with different technologies (**Table S19**). In most cases, comparisons made across short and long read datasets were consistent with results from comparisons made within read types for each pipeline.

When a 99% ANI threshold was used to group strains, Graphtyper with either short or 400 bp fragmented long reads was able to correctly genotype 19 of 20 truth-set genotypes (**Tables S15, S18**). In this analysis, the final pair, strains L98/93 and L219/93, had >50,000 SNPs to the chosen reference genome and were inferred to differ by ∼27 SNPs (**Table S15**, group 723). Clair3 with full-length reads or short reads and a quality filter of QUAL > 28 correctly genotyped 17 and 16 of 20 truth-set genotypes, respectively (**Tables S16, S18**).

### Epidemiology of Plant Clinic strains

Once we had identified an optimal pipeline and strategy for variant calling, we characterized the identity and inferred transmission patterns for the newly sequenced strains. The 116 sequenced strains are genetically diverse and represent each of the major lineages of agrobacteria (**Figure S3**). Many strains are closely related to previously sequenced agrobacteria from other hosts or locations. Most strains carry oncogenic plasmids, suggesting that they are likely to be pathogenic. Genotypes inferred from the Graphtyper analysis of strains partitioned using a 99% ANI threshold were visualized as minimum spanning networks (**Figure 4; Table S15**). Both short reads and 400 bp long-read fragments were consistent in identifying epidemiological links between or within locations. Strains CG163 and CG165 belong to the same genotype but were collected from two different vineyards. These data suggest a potential transmission event or acquisition from a common source. Likewise, two non-pathogenic strains (CG159, CG160) sampled from grapevine in California in 2000 are genetically identical to a non-pathogen strain also sampled from grapevine in Oregon in 2015 (15-2141-1B). Three strains (CG159, CG160, and CG161) collected from nursery N25 in 2000 represent multiple genotypes (**Figure 4**). Likewise, several genotypes of strains isolated from nursery N7 in 2022 differ by more than 30,000 SNPs (**Figure 4**). Both examples represent co-occurring infections at a single location.

**Figure 4:**
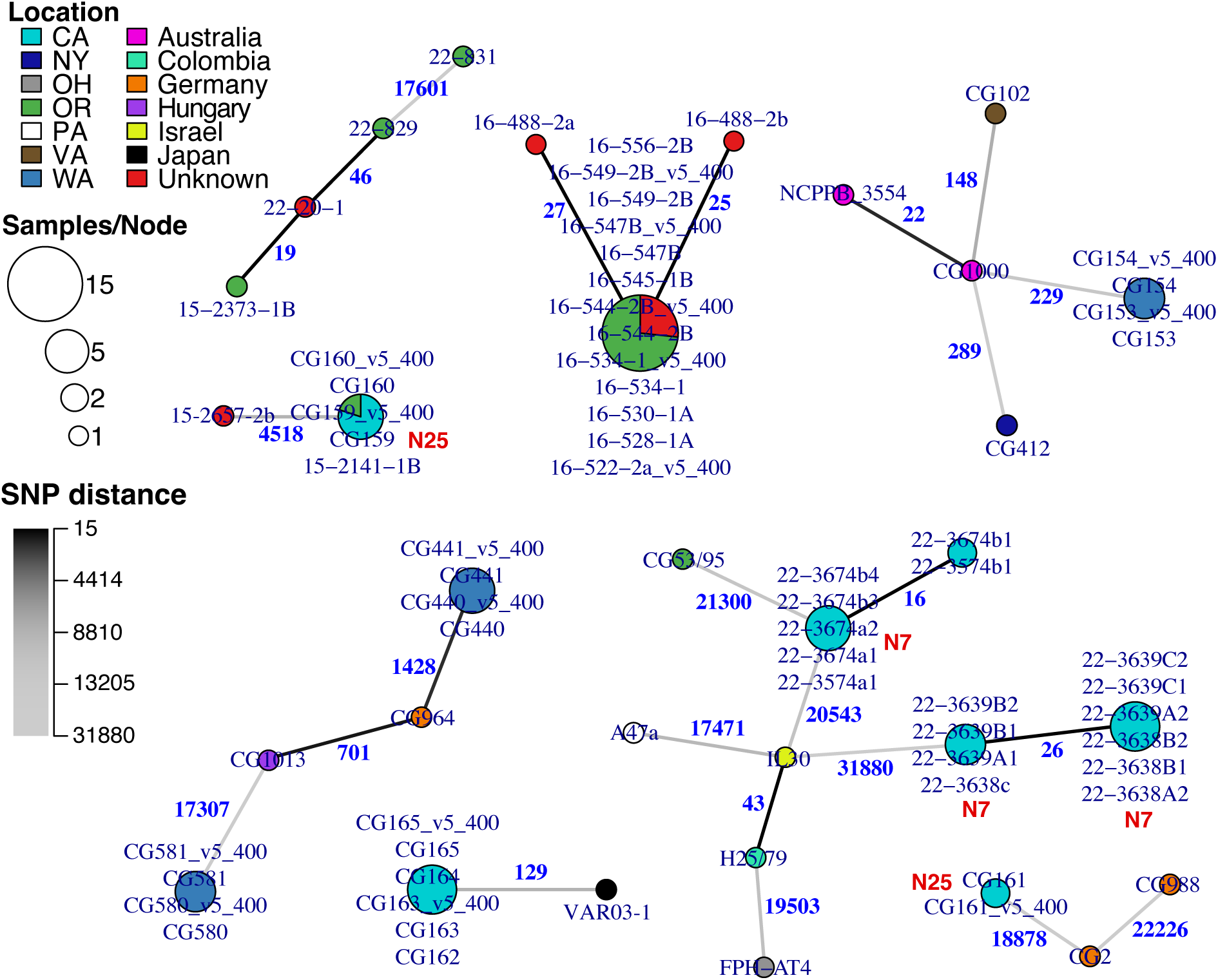
Whole genome sequencing data reveals epidemiological patterns. Minimum spanning networks representing relationships between select strains. Nodes represent whole-genome genotypes inferred from strains with ≤15 SNP differences. Edge labels and color represent the number of SNP differences between genotypes. Node size indicates the number of strains of each genotype, the proportion of each color indicates the location of isolation (country or U.S. state) for strains of that genotype. Genotypes from select nurseries are labelled in red text. Strains were analyzed in groups with >99% ANI similarity. Genotypes were inferred from variant calls produced by Graphtyper with either short or 400 bp long read data (labeled “_v5_400”).

## Discussion

These results and other recent studies suggest that Nanopore long reads are now of high enough quality to be used for genome assembly and variant calling on their own (8, 34–37). With the latest basecalling models and sufficient depth of sequencing, long reads can be used to assemble complete genomes with very few errors. The most serious errors are indel errors, as these can cause apparent frameshift mutations that disrupt gene annotation. However, these errors are also reduced in reads produced by the latest basecalling models (**Table S4**). While long-read assemblies are likely of acceptable quality on their own, shallow Illumina sequencing can also be added to ensure perfect assemblies (12). However, for most use cases this may not be necessary. Long-read assemblies are sufficient for identifying an organism, inferring phylogenetic relationships, and characterizing the presence of virulence genes or genes involved in antimicrobial resistance (AMR).

Our variant calling results are comparable to previous studies, but we identified different pipelines as most accurate (34, 35, 37). These differences are likely due to several factors, including the inclusion of different pipelines, such as Graphtyper and GATK. Previous studies used closely related reference genomes (34, 35, 37). In this study, we tested datasets representing population-level studies rather than a direct comparison of strains known to be nearly identical. Other studies introduced simulated changes in the genome of sequenced strains to use as truth-set variants (37). In this study, strains were sequenced with both Illumina and Nanopore technologies, and variants produced from short reads were used as the truth set. It is possible that long reads may identify additional true variants that are not captured by short reads, particularly in repetitive regions (37). To avoid this, we identified pairs of strains that are identical in direct comparisons with short and long reads. These strain-pair genotypes served as an additional truth set that is robust to this potential issue. Tested pipelines with the greatest precision and recall also recovered the greatest number of known strain-pair genotypes in population-level analyses. Pipelines that use pangenome graphs, such as Graphtyper, had the greatest consistency in calling variants across strains when using a distant reference genome (16).

Results from this study show that fragmenting Nanopore reads into shorter sequences is important for ensuring accurate variant calling with pipelines designed for short reads. All short-read pipelines produced more accurate variant calls when long reads were first fragmented into shorter sequences (**Figure 3**). Once long reads were fragmented, pipelines designed for short reads were as or more accurate than tools designed for long reads. The reduced accuracy with full-length reads is not likely due to poor mapping of full-length reads to a distant reference, as mapping quality statistics for full-length long reads were comparable to fragmented reads (**Figure S5; Table S11**). Importantly, fragmenting mapped full-length reads “in place” resulted in nearly identical variant calls as for pre-fragmented reads (**Table S10**). These “in-place” fragmented read alignments have identical reference coverage and read depth as the full-length read alignments, yet variant calls were much improved. This suggests that these variant calling differences are likely due to an inability of pipelines designed for short reads to correctly analyze long reads. Modifying these tools to support long reads would allow users to use the most accurate tools without losing the information present in full-length reads. For now, fragmentation can be performed prior to or after read mapping to promote accurate variant calling.

Previous studies found the long-read pipeline Clair3 to be the most accurate variant caller with long reads (37). For our datasets, Clair3 was among the top variant calling tools, but only when a more stringent variant filtration threshold was used (**Figure 2B; Table S7**). Using Clair3 with the developer-recommended filtering threshold of QUAL > 2 resulted in a larger number of false positive variants and the recovery of fewer truth-set genotypes (**Figures 2C, 3; Table S18**). This was especially the case when variants were called to a distant reference. Once a more stringent filtering threshold of QUAL > 28 was used, Clair3 with full-length reads was comparable in accuracy to short-read pipelines with fragmented long reads. However, variant calls produced by Clair3 from short reads recovered few truth-set genotypes at either filtering threshold (**Table S18**).

Most variant calling tools are only designed for either short or long reads. Few tools explicitly have options for mixed datasets. This results in compromises in choosing parameters or settings for each run. For example, using long reads with Graphtyper requires setting an option that ignores filtering by read pair mapping (“--no_filter_on_proper_pairs”). If a Graphtyper analysis has strains with paired short reads and other strains with long reads, read pairing will be ignored for the short-read datasets. This parameter did not dramatically impact variant calling with short reads in our analyses; however, it may be a problem for other datasets. Pipelines that include per-sample options for either short or long reads could improve results for mixed datasets. Alternatively, users should use only one read type or the other.

Caution should be taken in interpreting variant calls and genotypes inferred from mixed datasets of short and long reads. In some cases, comparisons across data types did not correctly infer a genotype for pairs of strains, but comparisons within data types did. For example, with Graphtyper and 95% ANI groups, analyses within 400 bp long-read or short-read data for strains CG153 and CG154 inferred them to be the same genotype (**Tables S10, S18**). But, when comparing across technologies, the 400 bp long-read data of CG154 were not genotyped with the short-read data of strain CG153 (**Tables S10, S19**). However, by grouping strains with a 99% ANI threshold, we eliminated most issues with genotyping across read types. In this analysis, short and long read data correctly grouped CG153 with CG154 in all comparisons (**Figure 4; Table S19**). We recommend using Graphtyper with fragmented long-reads for characterizing mixed datasets. Clair3 with more stringent filtering is also accurate when only long-read data are used.

Using a higher ANI threshold or identifying sub-lineages to select closer reference genomes may improve variant calling and genotyping (32, 38). Even at a 99% ANI threshold, strains can differ by >40,000 SNPs. We observed several cases where a 99% ANI threshold is still too distant for accurate variant calling. For example, using strain CA75/95 as reference for 99% ANI group 723, the short and long reads of strains L219/93 and L98/93 did not genotype together (**Table S18**). However, when variants were instead called using L219/93 as reference, the read sets and strains were genotyped together as expected (**Table S18**). If strains appear to be closely related but not the same genotype, we recommend re-analyzing with a reference genome more closely related to those strains.

Generating whole genome data is now easier and less expensive than before. Individual labs or plant clinics can prepare libraries and perform Illumina short read or Oxford Nanopore long read sequencing in the lab with relative ease. These results suggest that either technology is sufficient for genome assembly and variant calling in most cases. Both Illumina and Oxford Nanopore have released DNA sequencers that can be purchased by individual labs. An advantage of long read sequencing is that libraries are generally less time consuming and easier to prepare in the lab. If sequencing is performed in house, costs can be cheaper, and results acquired more quickly, than with short read sequencing. Rapid advances in sequencing technology provide a major opportunity for plant clinics and labs to generate data with the highest resolution for diagnostics and epidemiology.

## Materials and Methods

### Genome sequencing and analysis

Strains were selected from the Larry Moore culture collection or from those collected by the OSU Plant Clinic (**Table S1**). Additional strains were shared by the California Department of Food and Agriculture (CDFA). Strains were cultured in MGYs liquid media or plates at 28°C (39). The Promega Wizard Genomic DNA Extraction kit (Promega; Madison, WI) was used to extract DNA from cell pellets of each strain. The protocol was modified to include an overnight lysis step at 37°C and mixing by inverting the tube rather than pipetting. All strains were sequenced with both Illumina and Oxford Nanopore technologies. The SeqWell PurePlex kit was used to prepare libraries for Illumina sequencing. Illumina libraries were sequenced (2×150 bp) on an Illumina MiniSeq (Illumina; San Diego, California) in the OSU Plant Clinic. The Rapid Barcoding 96 V14 kit (SQK-RBK114.96) was used to prepare multiplexed libraries for Nanopore sequencing. Nanopore libraries were sequenced on an R10.4.1 PromethION flow cell on a P2 solo sequencer (Oxford Nanopore; Oxford, UK). Dorado v.0.5.0 with the default parameters, SUP basecalling, and different basecalling models was used to basecall nanopore reads (40).

FastQC was used to check Illumina reads for quality (41). NanoPlot v1.42.0 with the default parameters was used to check Nanopore reads for quality (42). Spades v3.15.3 with the parameters “--phred-offset 33 –careful -k 21,33,55,77,99” was used to assemble genomes from only Illumina reads (20). For all assemblies, contigs with <500 bp in length or <5X coverage were removed. Flye v2.9.2-b1786 with the parameters “--scaffold --read-error 0.03 was used to assemble genomes from only Nanopore reads (19). Medaka v.2.0.0 with the model r1041_e82_400bps_sup_v4.2.0 or r1041_e82_400bps_sup_v5.0.0 was used to polish Flye assemblies (23). Polypolish v0.5.0 with the default parameters was used to polish Flye assemblies with Illumina reads (11). Unicycler v0.5.0 with the parameter “--mode normal” was used to create a hybrid assembly from both Illumina and Nanopore reads (21). Beav v0.2 with the parameter “--agrobacterium” and Bakta v.1.8.2 was used to annotate genomes (43, 44). BUSCO with the parameters “-m genome −l rhizobium-agrobacterium_group_odb10”, Quast v5.0.0 with the default parameters, and the Samtools v.1.19.2 subcommand “depth” were used to gather genome assembly statistics (15, 45, 46). MEME v5.4.1 with the parameters “-dna -nmotifs 10” was used to identify motifs around putative Flye assembly errors identified by Polypolish polishing (47). Motifs were filtered to those with an evalue < 1. Bandage v.0.8.1 with the default parameters was used to visualize assembly graphs (48). FastANI v1.1 with the default parameters was used to calculate pairwise average nucleotide identity (ANI) and cluster strains into groups based on 95% or 99% thresholds (49).

The assembly of strain CG160 was found to have an unusual number of errors. BBTools sendsketch.sh v39.06 with the default parameters was used to identify contaminants (50). Sourmash v4.8.11 sketch and compare were used to generate and compare signatures for the contaminated plasmid and the plasmids of a closely related strain (51). BBTools seal.sh v39.06 with the parameters “pattern=out_%.fq ambig=first” and combined reference assemblies were used to filter out contaminated reads (50). The assemblies of *Rhizobium rhizogenes* K84 (NCBI: GCF_000016265.1), *Rhizobium tumorigenes* B21/90 (NCBI: GCF_019355815.1), CG160 without the contaminated plasmid, and the equivalent plasmid from closely related strain CG159 were used as input to partition reads. Non-contaminant reads mapping to the CG160 assembly and CG159 plasmid were combined and used in the previously described analyses.

### Phylogenetic analysis

NCBI Datasets was used to download all publicly available genomes of the *Agrobacterium*/*Rhizobium* complex from NCBI on July 19^th^, 2023 (52). Automlsa2 v0.8.1 with the parameters “--allow_missing 4 --missing_check” was used to generate a maximum likelihood multi-locus sequence analysis (MLSA) phylogeny including newly sequenced and NCBI strains (53, 54). The protein sequences AcnA, AroB, CgtA, CtaE, DnaK, GlgB1, GlyS, HemF, LeuS, LysC, MurC, PrfC, RecR, RplB, RpoB, RpoC, SecA, SMc00019, SMc00714, SMc01147, SMc01148, SMc02059, SMc02478, ThrA, and TruA from *Sinorhizobium meliloti* strain 1021 were used as references (55). IQ-TREE 2 v2.2.0 with parameters “-m MFP -B 1000 -alrt 1000 --msub nuclear --merge rclusterf” was used to generate a maximum likelihood phylogeny (56). The R package ggtree was used to plot phylogenies (57).

### Whole genome SNP calling and comparisons

Whole genome SNP analyses were performed using Illumina short reads, full-length Nanopore reads, or fragmented Nanopore reads. All newly sequenced strains, including those with short reads or both short and long reads, and strains downloaded from NCBI were analyzed together. Seqkit v0.16.1 sliding with parameters “-W 150 -s 150” or “-W 400 -s 400” was used to split Nanopore reads into 150 bp and 400 bp fragments, respectively (58). Species groups were determined by single-linkage clustering of ANI values of short-only, hybrid, or NCBI assemblies, with minimum ANI thresholds of 95% or 99% (49). Within species groups/lineages, references were selected among hybrid or NCBI assemblies based on assembly completeness, as determined by a low number of contigs and a high N50. Kingfisher v0.2.2 subcommand “get” and parameters “-m ena-ascp aws-http prefetch” was used to download Illumina reads for NCBI strains (59). Bwa v0.7.17-r1188 mem with the parameter “-M” was used to map Illumina reads to a reference within each group (60). Minimap2 v2.26-r1175 with parameter “-ax map-ont” was used to map Nanopore reads to a reference within each group (61). Samtools v1.19.2 with the subcommand view and parameter “-F 256” was used to keep only primary alignments for mapped long reads (15). For the fragmented in-place analysis, a custom python script was used to fragment mapped long reads “in-place” into 400 bp sequences, maintaining all mapping information and quality from the original alignment.

The developer-suggested parameters and filtering thresholds were used for each tested pipeline. GATK v4.6.2.0 HaplotypeCaller with the parameters “-ERC GVCF -ploidy 1 - allowPotentiallyMisencodedQuals” was used to call variants for each dataset (17). GATK CombineGVCFs and GenotypeGVCFs with the default parameters were used to combine partial variant calls across strains. GATK SelectVariants with parameter “-selectType SNP” was used to select SNP variants only. GATK VariantFiltration with parameter “--filterExpression QD < 2.0 SOR > 3.0 QUAL < 30.0 FS > 60.0 MQ < 40.0 MQRankSum < −12.5 ReadPosRankSum −8.0” was used to hard filter SNPs. The GATK tool SelectVariants was used with parameter “--excludefiltered” to remove filtered SNPs. Graphtyper v2.7.3 with the parameter “--no_filter_on_proper_pairs” was used to call SNPs (16). The GATK VariantFiltration tool with parameters “-G-filter ‘isHet == 1’ -g-filter-name ‘isHetFilter’ --set-filtered-genotype-to-no-call’” was used to filter heterozygous calls from Graphtyper vcf files. The vcflib program vcffilter with parameters “-f ‘ABHet < 0.0 | ABHet > 0.33’ -f ‘ABHom < 0.0 | ABHom > 0.97’ -f ‘MaxAASR > 0.4’ -f ‘MQ > 30’” was used to filter Graphtyper variant calls for quality (62). Bcftools v1.17 mpileup with parameters “-x -I -Q 13 -h 100 -M 10000 -a ‘INFO/SCR,FORMAT/SP,INFO/ADR,INFO/ADF”, subcommand call with parameters “-m -ploidy 1 -V indels”, and the apply_filters.py script with parameters “-q 27 -M 55 -V 0.00001 -x 0.2 -K 0.9 -s 1 -d 5” were used to call SNPs and filter for quality (15, 36). Clair3 v.1.0.4 with parameters “--no_phasing_for_fa --include_all_ctgs --haploid_precise –gvcf” and either “--platform=’ont’ -- model_path=“${CONDA_PREFIX}/bin/models/r1041_e82_400bps_sup_v500”” or “--platform=’ilmn’ --model_path=”${CONDA_PREFIX}/bin/models/ilmn”” for Nanopore or Illumina reads, respectively, were used call SNPs (18). Bcftools merge with the parameter “-0” was used to merge Clair3 gvcf files. The vcflib program vcffilter with parameters “-f ‘QUAL > 2’” or “-f ‘QUAL > 28’” was used to filter Clair3 variant calls for quality. The R package poppr v.2.9.5 was used to calculate pairwise SNP distances and plot minimum spanning networks (33). Bcftools view with the parameter “--samples” was used to extract individual strain SNP calls for both short reads and long reads fragmented into 400 bp. These files were filtered to only contain non-reference SNP calls.

To identify truth-set genotypes, short or full-length long reads of each strain were mapped individually to the hybrid assembly of all other strains, along with reads of that reference, and used to call variants. Bwa and Graphtyper with the above-described parameters were used to align and call variants with short reads (16, 60). Similarly, minimap2 and Clair3 with above-described parameters were used to map full-length long reads and call variants for the same comparisons (18, 61). Variant calls were filtered as described above. Strain pairs where the short or long reads of both strains were inferred to have fewer than 15 pairwise differences were considered to be a genotype.

Variant calling accuracy and consistency across technologies was then characterized for the 35 strains with both Illumina and Nanopore reads from the above-described analyses. VCFCompare v.1.0 was used to calculate overall precision and accuracy scores (63). Vcfdist v2.5.2 with the parameters “--max-qual 60 --min-qual 0 --cluster gap 50” and Illumina SNPs as the truth dataset was used to calculate accuracy scores for Nanopore SNPs across quality thresholds (64). For precision vs. recall curves, true positives, true negatives, false positives, and false negatives were each summed across the 35 strains at each quality score threshold for calculation of precision and recall. The R package FSA was used to perform the Kruskal-Wallis test followed by a Dunn’s test with Holm Method *p*-value adjustment to assess statistical significance of precision and recall between pipelines (65).

## Acknowledgements

We would like to thank the California Department for Food and Agriculture (CDFA) for sharing *Agrobacterium* strains. We thank the Department of Botany and Plant Pathology for supporting the CQLS high performance computing infrastructure. This work was supported by startup funds from the Department of Botany and Plant Pathology at Oregon State University. This project was funded in part by USDA NIFA through the Western Integrated Pest Management Center, award SA 18-4060-41, and by the National Institute of General Medical Sciences of the National Institutes of Health under award number 1R35GM160469.

## Data Availability

Final genome assemblies and raw read data were uploaded to NCBI under BioProject PRJNA1255661. All genome assemblies and scripts for this project can be found in the following GitHub repository: https://github.com/weisberglab/nanopore_quality_manuscript. Variant calling vcf files for each dataset are also available in the following Zenodo repository: https://doi.org/10.5281/zenodo.15652634.

