## Supplemental Figures for "A comparison of short- and long-read whole genome sequencing for microbial pathogen epidemiology"

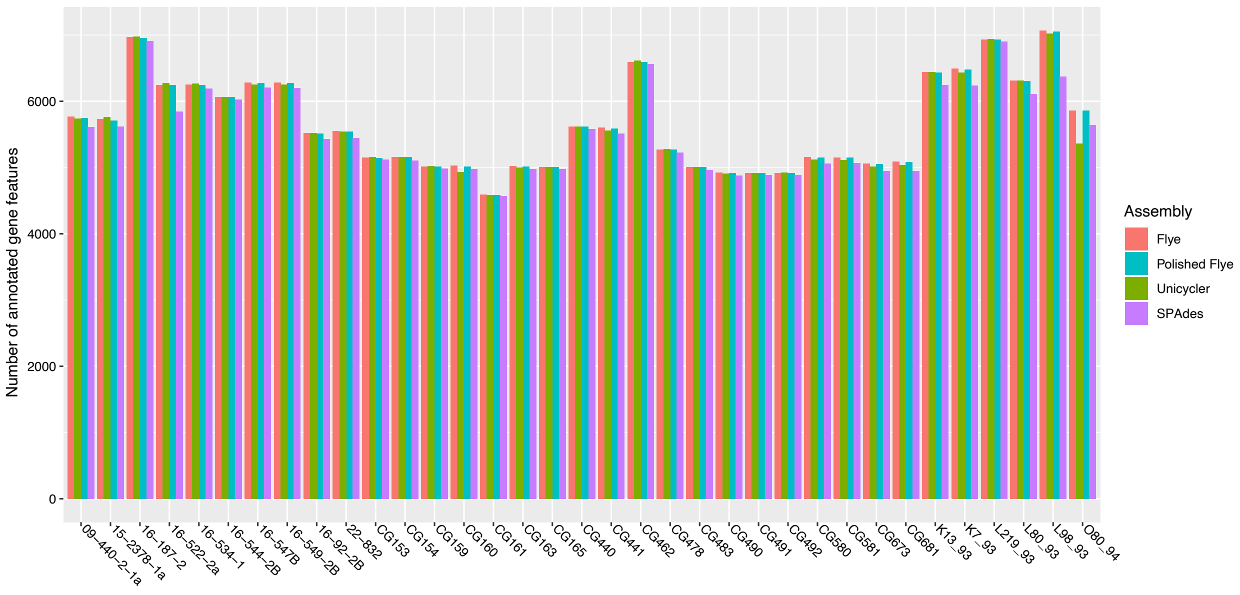


**Figure S1: Total number of genes annotated in each assembly.** Bars indicate the number of genome features annotated in short-read polished and unpolished Flye long-read assemblies, Unicycler hybrid assemblies, and SPAdes short-read assemblies. Assemblies are grouped by strain, bars represent the count of genome features. Colors represent different assembly tools or datasets.

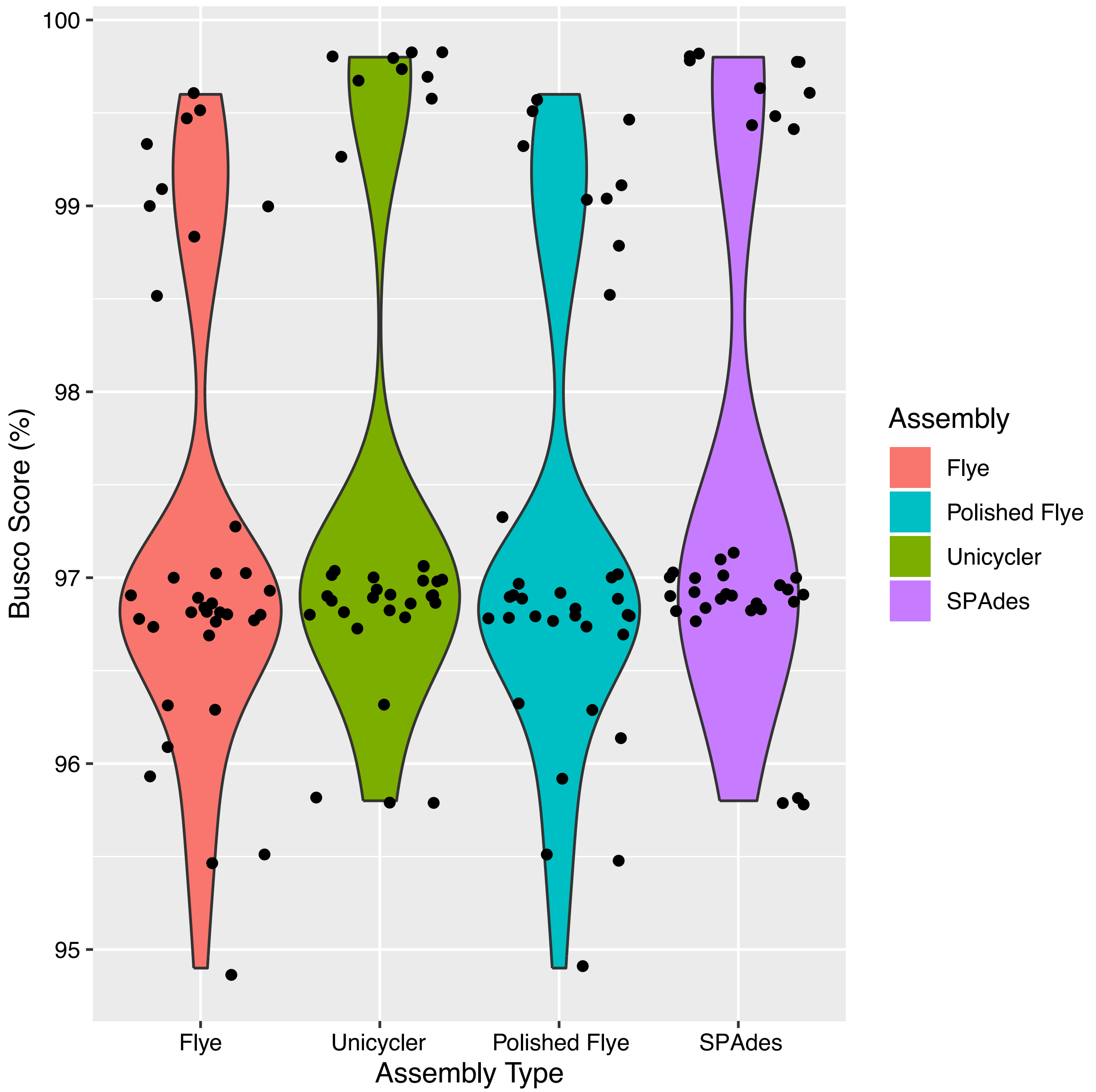


**Figure S2: BUSCO score comparison across assembly methods and datasets.** BUSCO scores for each individual strain are plotted as points within each associated assembly type. The y axis represents the percentage of housekeeping genes found in each strain based on the *Agrobacterium/Rhizobium* BUSCO database.

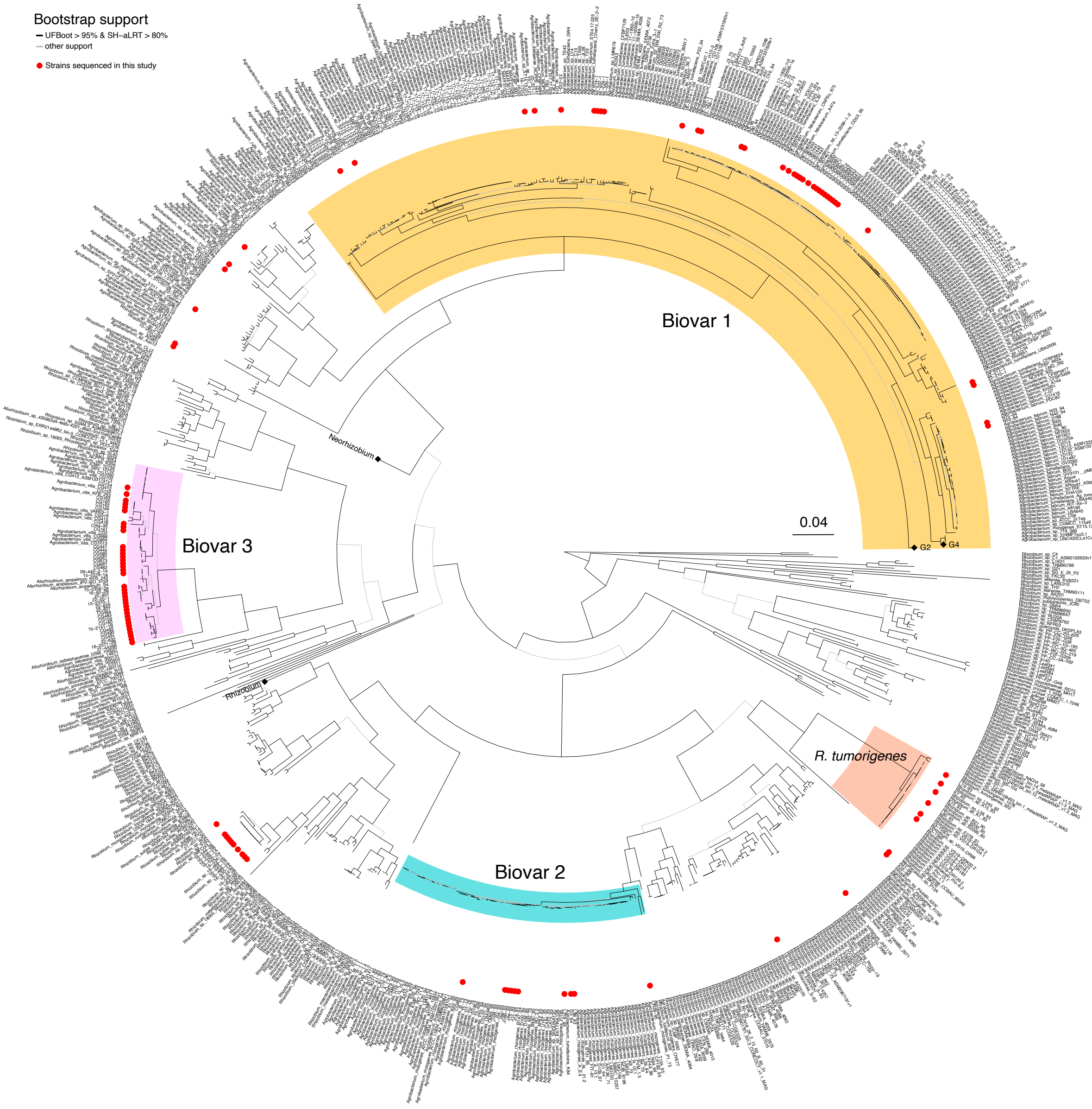


**Figure S3: Agrobacteria sequenced at the OSU Plant Clinic are genetically diverse.** Multi-locus sequence analysis (MLSA) phylogeny of the agrobacteria/rhizobia complex. Strains sequenced as part of this study are indicated by a red point. Major agrobacteria lineages (biovars) are shaded. Biovar 1 is highlighted yellow, Biovar 2 is highlighted blue, *R. tumorigenes* is highlighted salmon, and Biovar 3 is highlighted pink. Collapsed branches are indicated by diamonds and labeled. The tree is midpoint rooted. Branch color indicates bootstrap support. Branches with ultrafast bootstrap (UFboot) >= 95% and SH-aLRT >= 80% are colored black, other branches are gray. Branch scale is indicated by a bar.

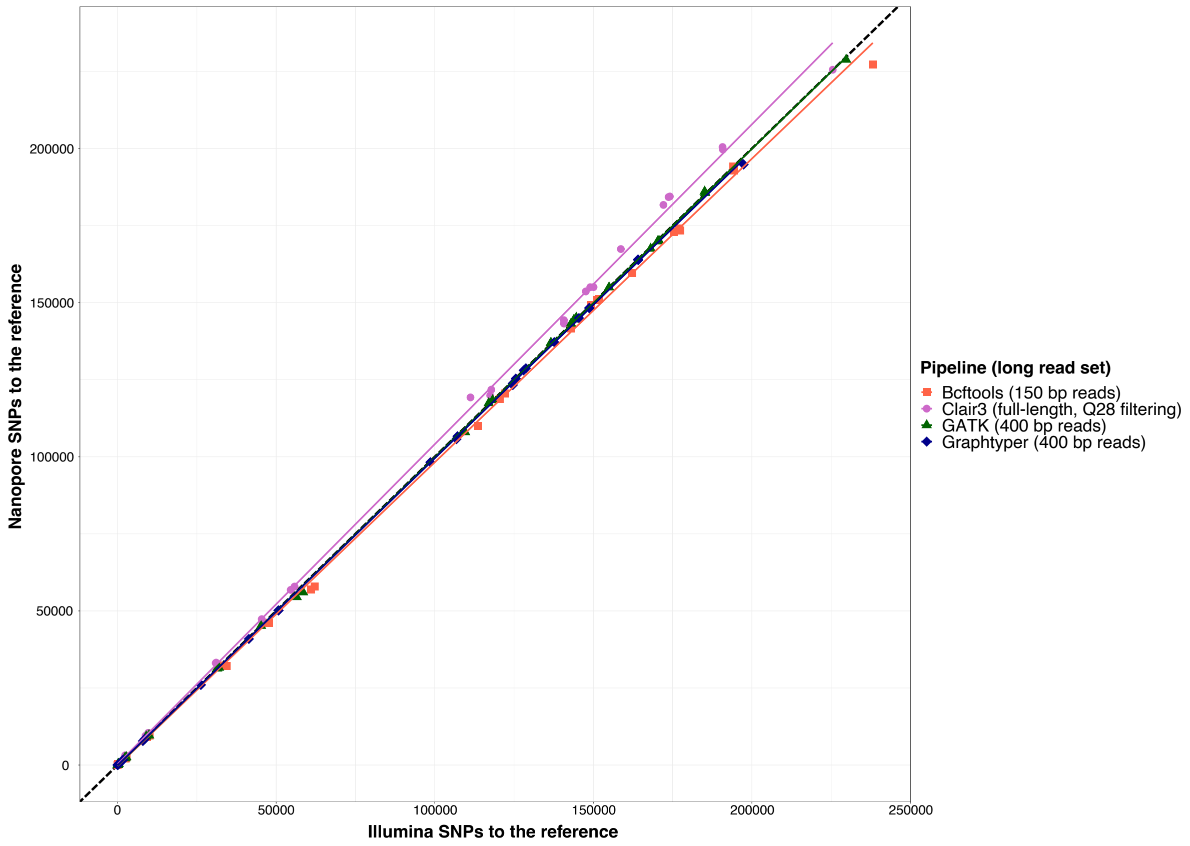


**Figure S4: Variant calling consistency among short and long-read datasets for each pipeline.** The long-read dataset with the greatest accuracy for each pipeline is shown. Each strain is plotted as a point with the number of SNPs to the reference called using Illumina reads on the x axis and SNPs called using long-read datasets, either full-length or fragmented, on the y axis for each variant calling pipeline. Best fit lines are colored by pipeline. The dashed black line represents a predicted 1:1 ratio between SNPs to the reference called using short and long reads.

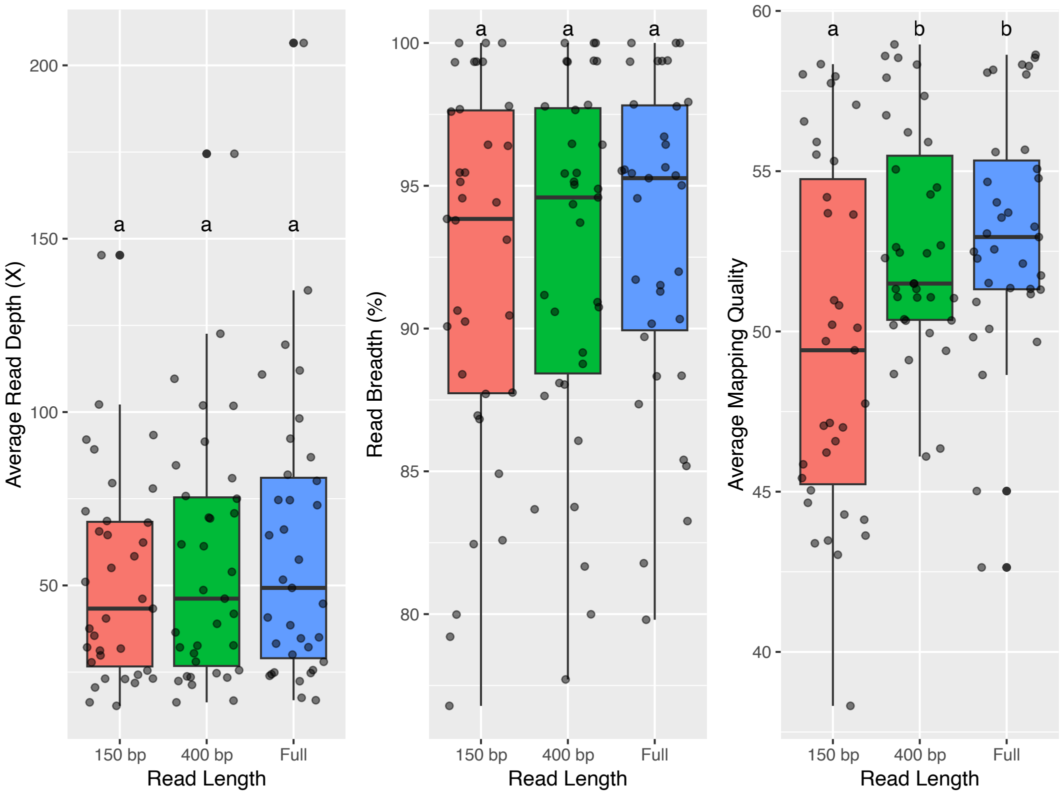


**Figure S5: Read mapping metrics comparison for different long-read fragment lengths.** Read mapping metrics from individual strains are plotted as a point for each associated read length and summarized as boxplots. Average read depth represents the number of reads per base, read breadth represents the percentage of the reference genome covered by reads, and average mapping quality uses Phred scoring. Kruskall-Wallis test *p*-values for read depth, read breadth, and mapping quality by read length are 1, 0.74, and 0.03. Letters above bar plots indicate multiple comparison group from Dunn's post-hoc testing.

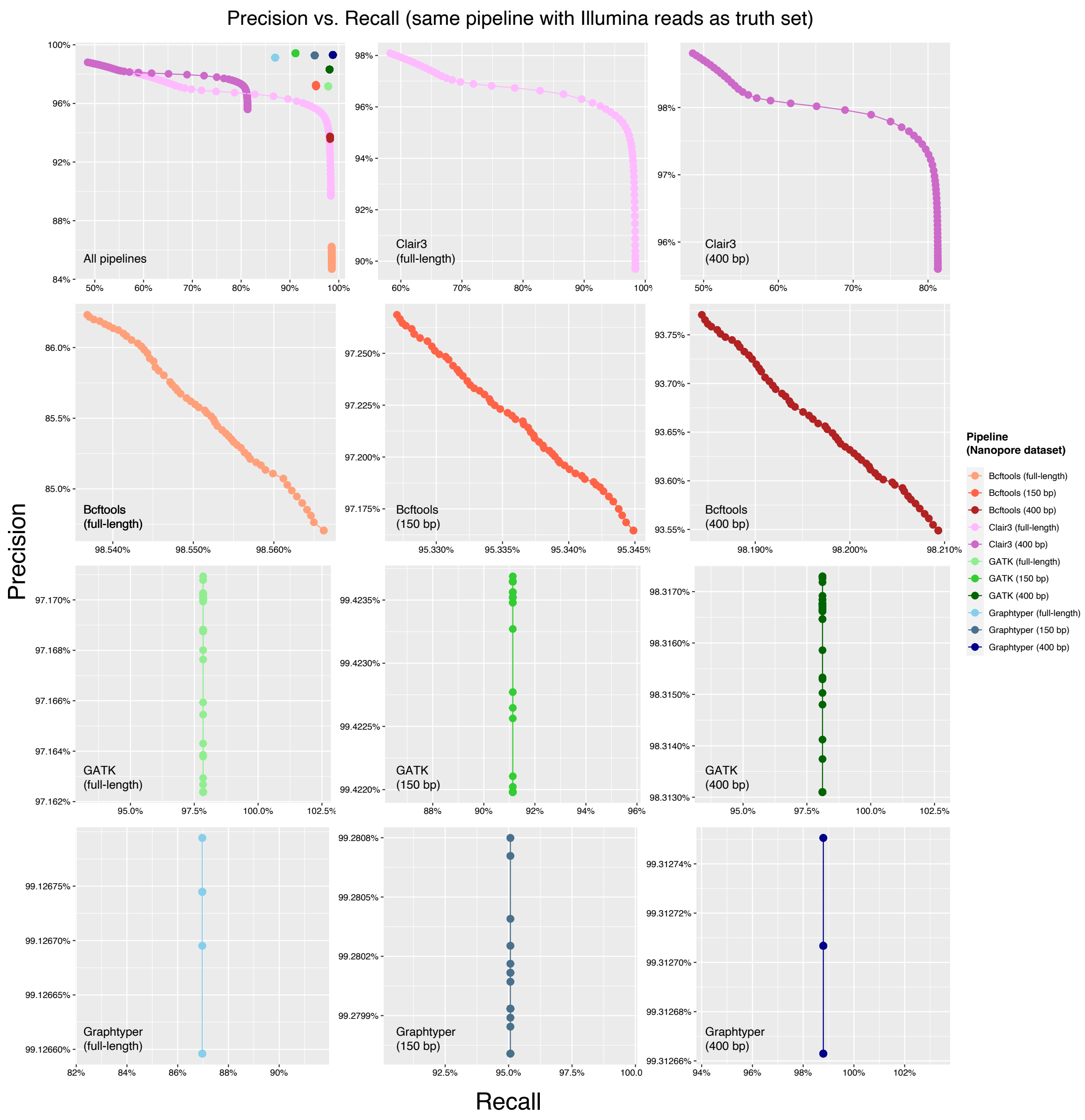


**Figure S6: Precision vs. recall plots for variant calling pipelines with long-read datasets (same pipeline truth).** Precision and recall scores were calculated for variant calls produced by each pipeline using full-length or 400 bp long reads at quality score (QUAL) thresholds from 0 to 60. Truth sets were variant calls produced by the tested pipeline with Illumina short reads. Precision and recall scores are represented as percentages. Scores calculated using variants across all strains at a given quality threshold are represented as points and summarized as lines. The upper left figure shows precision vs. recall for all pipelines and read sets, other figures show results for individual pipelines and read sets.

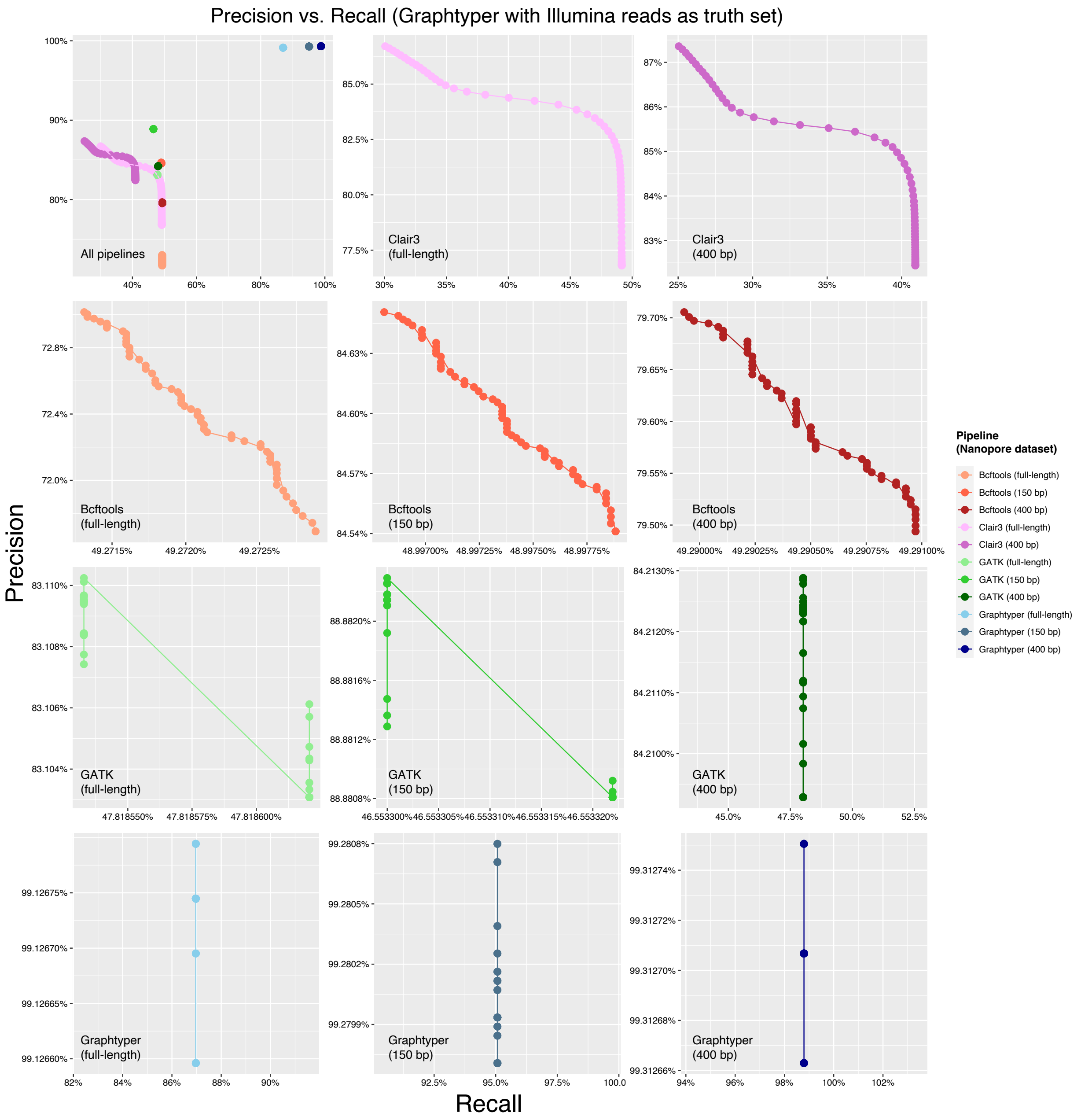


**Figure S7: Precision vs. recall plots for variant calling pipelines and read datasets (Graphtyper truth).** Precision and recall scores were calculated for variant calls produced by each pipeline using full-length or 400 bp long reads at quality score (QUAL) thresholds from 0 to 60. Truth sets were variant calls produced by Graphtyper with Illumina short reads. Precision and recall scores are represented as percentages. Scores calculated using variants across all strains at a given quality threshold are represented as points and summarized as lines. The upper left figure shows precision vs. recall for all pipelines and read sets, other figures show results for individual pipelines and read sets.
